# Long-term iododeoxyuridine labelling analysis finds no evidence of ovarian germline stem cell activity in adult or ageing mice

**DOI:** 10.1101/2025.04.01.646696

**Authors:** Chinelo Miriam Onochie, Ahlam Saleh Alamri, Kaja Kostanjevec, Alex Francesca Brown, Samuel Luke Burton, Sandra Susi Koigi, Stephanie Lee Wattie, Jon Martin Collinson

**Affiliations:** School of Medicine, Medical Sciences and Nutrition, University of Aberdeen, Institute of Medical Sciences, Foresterhill, Aberdeen AB25 2ZD, UK

**Keywords:** Germline, stem cells, oocytes, gene expression, cell division, immunochemistry, histology, VASA

## Abstract

The purpose of this study was to determine whether ovarian germline stem cells or any adult mitotic cell type contribute to the adult oocyte pool in vivo from early adult through to aging mice. Long-term thymidine analogue labelling assays using iododeoxyuridine (IdU) were performed to identify mitotic cells and their descendants by immunohistochemistry for IdU and germline markers Ddx4 and Oct4. C57BL/6 mice at 10 weeks, 5 months and 12 months old were given IdU for 21-30 days through their drinking water to label all dividing cells, followed by washout periods of 0-10 weeks. Over 60,000 ovarian somatic cells and oocytes were scored in 111 ovaries. No double labelling of IdU with germline stem cell markers was found in oocytes at any stage of maturity, in mice at any age. In contrast, IdU exposure during embryogenesis resulted in large numbers of labelled oocytes in postnatal mice, as expected. We were therefore unable to confirm mitotic activity in stem cells or any other progenitor replenishing the oocyte pool in C57BL/6 mice. There is no evidence that quiescent stem cells are activated when the oocyte pool is depleted by age.

## INTRODUCTION

A central dogma of reproductive biology is that female mammals acquire a fixed reserve of oocytes before birth which depletes from puberty to menopause without replacement (Zuckerman, 1951). The ovarian reserve reflects primarily the number of primordial (non-growing) follicles, together with follicles recruited into the later pre-antral and antral stages of development ultimately capable of ovulation. During development, primordial germ cells (PGCs) migrate to the genital ridges and increase in number through mitosis. Eventually these PGCs, now called oogonia, enter meiosis and become integrated into primordial follicles where their development is arrested during the first meiotic division, after DNA replication. The long-standing view is that meiotic arrest fixes the ovarian reserve at this stage of development and that neo-oogenesis does not occur in adult life (Grive and Freiman, 2015).

Johnson et al. (2004) challenged the central dogma; they argued that, in mice, oocyte numbers at birth were likely insufficient to last the reproductive life of the individual. They presented evidence that the oocyte pool was replenished by a population of mitotic stem cells. The finding was controversial, however multiple groups have since published in vitro or in vivo data suggesting that the oocyte pool can be replenished or modulated in adult life, possibly by regenerative stem cells in humans and mice (Kerr et al., 2006; Pacchiarotti et al., 2010; Tilly and Telfer, 2009; Virant-Klun, 2015; White et al., 2012; Niikura et al., 2009; Niikura et al., 2010; Wang et al., 2017; Zou et al., 2009; Zou et al., 2011). In contrast, there is also a significant body of data showing, through lineage analysis, gene expression and transplantation experiments, that neo-oogenesis *does not* occur in mammals (Begum et al., 2008; Kerr et al., 2012; Lui et al., 2007; Zhang et al., 2012; Zhang et al., 2015). A consensus point of view is that it may be possible to isolate or identify cells capable of regenerating oocytes in vitro or in some experimental situations. The objective of this study was to further determine whether stem cells, or any other mitotic cells in mouse adult tissues exist or not, using long term labelling of female mice with the thymidine analogue 5-iodo-2’-deoxyuridine (IdU) that can be incorporated into replicating DNA and subsequently detected by immunohistochemistry. In addition to assaying young females that may not require stem cells, evidence for stem cell activity was sought in mice approaching the end of their peak reproductive potential (5 months old) and beyond normal breeding age (12 months old), on the basis that stem cell activity may activate as the oocyte pool depletes. If any mitotic germline stem cells exist in adult ovaries, long term IdU labelling should result in at least occasional detection of IdU-positive oocytes derived from those mitoses. In contrast if the long-standing dogma that the entire lifetime oocyte pool is set during fetal life and arrested during meiosis I, no cells in the adult oocyte lineage should ever replicate their DNA and will hence always be IdU-negative.

## METHODS

### Mice

All experiments were performed under the UK Animals (Scientific Procedures) Act 1986. Ethical permission was granted by the University of Aberdeen Ethical Review Committee. Mice were group-housed in individually vented cages with standard wood pellet, paper bedding and environmental enrichment. Food and water were provided *ad libitum*.

### Preparation of iododeoxyuridine drinking water

5-iodo-2’-deoxyuridine (I7125, Sigma) was dissolved 50 mg/ml in 0.2 M NaOH. Aliquots were stored at −20°C. IdU was diluted to 1 mg/ml in sterile H_2_0 immediately before giving to mice. IdU drinking water was protected from light and replaced every 2-3 days.

### IdU labelling

Fifty, 10-week old virgin C57BL/6 female mice were administered 1 mg/ml IdU via their drinking water for 30 days to label even very infrequently dividing cells. They were subject to increasing periods of washout −0 weeks (20 mice), 3 weeks (10 mice), 6 weeks (9 mice), and 10 weeks (11 mice) - during which they were given normal drinking water.

5 mice at 5 months old and 5 mice at 12 months old were given 1 mg/ml IdU for 21 days, followed by washout periods of 0-9 weeks. At the end of each washout period, some mice from the group were killed and ovaries dissected and fixed in 4 % paraformaldehyde in phosphate-buffer saline (PBS), pH 7.4 for 3 hours at 4°C. Tissues were washed 3 times in PBS for 25 minutes, in 0.9 % NaCl, 50 % and 70 % ethanol for 15 minutes each at room temperature, and stored in 70 % ethanol at 4°C.

For labelling during embryogenesis, wild-type pregnant C57BL/6 female mice, >20 g weight, received a single subcutaneous injection of 200 μl 2 mg/ml IdU in sterile saline to the nape of the neck at embryonic day 14.5.

#### Histology and immunohistochemistry

Both ovaries were dissected from killed mice and normally both were analysed unless one was lost or damaged during processing. Ovaries were dehydrated in 85 % ethanol for 20 minutes, 95 % ethanol 2x 20 minutes, 100 % ethanol 2x 30 minutes and finally once for 1 hour. Ovaries were cleared in xylene, 2x 5 minutes, and overnight before embedding in Paraplast Plus (Sigma P3683). 7 μm sections were cut and mounted on Poly-L-lysine coated glass slides. For immunohistochemistry, slides were dewaxed in Histoclear 2x 10 minutes and rehydrated through decreasing concentrations of ethanol into PBS. Antigen retrieval was achieved by boiling the slides in 0.1 M sodium citrate buffer (pH 6.0) for 20 minutes and washing for 5 minutes in PBS at room temperature. Blocking buffer (0.3 % BSA in PBS, 4 % donkey serum, 0.1 % Triton X-100) was added to each slide for 1 hour at room temperature. Primary antibodies: anti-IdU mouse monoclonal (Abcam: ab8955), anti-Ddx4 rabbit polyclonal (Abcam: ab13840) and anti-Oct4 rabbit polyclonal (Abcam: ab18976) were diluted 1:200 in blocking buffer and applied to each slide overnight at 4°C. The following day slides were washed 3x 5 minutes in PBS and a 1:200 dilution of secondary antibodies Alexa 488 goat anti-mouse (Molecular Probes: A21121) and Alexa 594 donkey anti-rabbit (Molecular Probes: A21207) in blocking buffer was applied for 2 hours at room temperature. Slides were washed 3x 5 minutes in PBS, and mounted with Vectashield with DAPI (Vector Laboratories, H-1200), for fluorescence microscopy analysis.

Negative control staining was performed by omitting primary antibody from a selection of slides at all ages.

#### Fluorescence Microscopy Analysis

Slides were examined on a Nikon E400 Eclipse fluorescence microscope. At least 10 randomly selected sections (non-consecutive, separated by at least 28 μm) from across each ovary were selected for analysis.

Cortical epithelial cells and oocytes in putative primodial, primary, secondary and fully mature (tertiary) follicles were counted, using DAPI and scored for IdU and Oct4 or Ddx4 labelling. Where a large oocyte was identified but the nucleus was not visible in the section, adjacent sections were examined such that as many mature oocytes were scored as could be found. Images were overlayed using Gimp 2.88 and Adobe Photoshop CS6.

## RESULTS

To confirm that thymidine analogue IdU labelling can successfully detect mitotic oogonia that go on to form primary oocytes, pregnant wild-type mice (n = 4) were given a single subcutaneous injection of IdU at E14.5 of pregnancy. Subsequent female pups were taken at 20 days (6 mice) or 45 days of age (7 mice). Ovaries were taken for immunohistochemistry for IdU and the oocyte/germline cell marker Ddx4. Ddx4 (Vasa/MVH) is a cytoplasmic ATP-dependent RNA helicase of the DEAD-box family that is involved in germ cell specification and is expressed in germ cell lineages in both sexes (Carrera et al., 2000; Erez, 2000; Woods and Tilly, 2013; Park and Tilly, 2015). As expected, all ovaries from all mice showed IdU-positive, Ddx4-positive primary oocytes, forming part of the ovarian reserve that was in mitosis at E14.5 (Figure 1).

**Figure 1.**
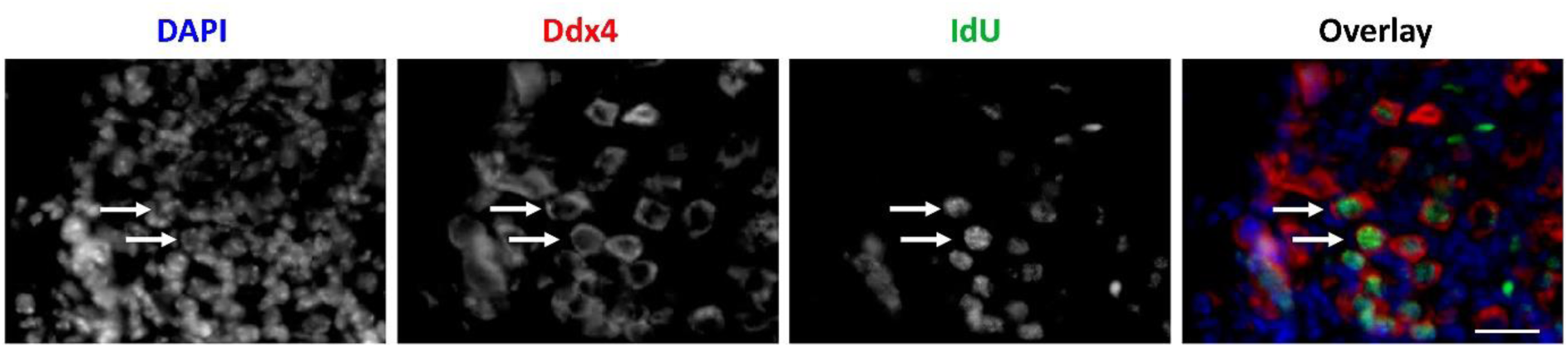
Thymidine analogue labelling of ovaries in utero. Pregnant mice were injected with IdU at E14.5 during the period when many primary oocytes are entering meiosis I and replicating DNA. Labelled pups were killed at postnatal day 45 for IdU (green) and Ddx4 (red) immunohistochemistry (n = 11). Each ovary contained many examples of IdU-labelled, Ddx4-positive oocytes. Representative section shown (left-right) of DAPI, Ddx4, IdU and Ddx4/IdU overlay, with some double-labelled cells indicated by arrows. Scale bar represent 10 μm.

### Analysis of mitotic activity in the germline of young adult female mice

To detect mitotic activity in the ovarian reserve of adult mice, females at 10 weeks - 12 months old were given drinking water containing IdU for 21 - 30 days, as described in Methods, to label all cells that replicated their DNA, which should include any active ovarian germline stem cells. Mice were killed after washout periods of 0, 3, 6, 9 or 10 weeks during which they received normal drinking water. Ovaries were taken for immunohistochemistry for IdU, Ddx4 and Oct4. Oct4 protein is a nuclear transcription factor present in germline cells that has an important role during oocyte growth (Shi and Jin, 2010; Bagheripour et al., 2017).

In young female mice (10 weeks old at start of experiment), many somatic cells of the ovary, including most of the ovarian cortex and epithelium, granulosa and theca cells were IdU positive after 30 days of labelling (Figure 2). With increasing periods of washout, IdU labelling was diluted out, first in rapidly dividing cells of the maturing follicles and more slowly in other tissues. At 10 weeks, most IdU was retained only in the cortex (Figure 2).

**Figure 2.**
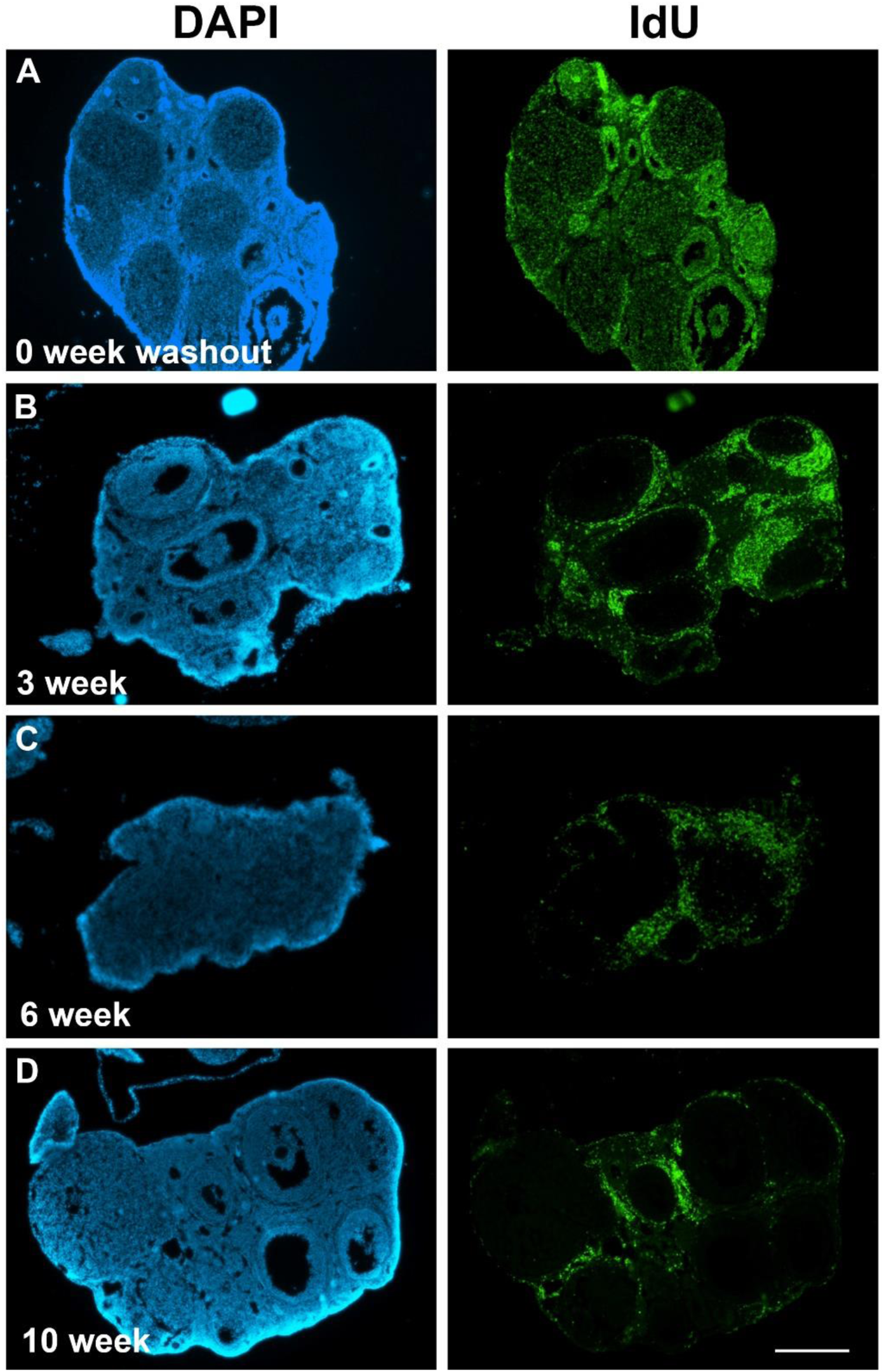
Labelling and retention of IdU (green) over a period of 10 weeks in mouse ovaries (A-D). Representative low-magnification images of tissue sections of ovaries from young adult mice labelled with IdU for 30 days, with 0, 3, 6 or 10-week washout. DAPI (blue) and anti-IdU (green). (A) At 0 week washout, immediately after labelling, widespread IdU accumulation was observed in cells in all somatic tissues including ovarian epithelium, ovarian cortex and granulosa and theca cells of maturing follicles. (B) At 3 weeks washout IdU retention was predominant but was rapidly diluted in actively dividing granulosa cells of maturing follicles. (C, D) At 6 and 10 weeks washout, IdU was further diluted and visible only in some cortical and epithelial cells. Scale bar represents 500 μm.

Immunohistochemistry with antibodies against Ddx4 or Oct4 revealed no double labelled cells (Figures 3; Table 1). For 82 ovaries across the four washout periods, over 6609 IdU-positive cells (of 58212 cells counted) were found around the periphery of the ovaries, in the epithelium and at the epithelial-cortical boundary, where many primordial follicles reside and represents a possible location for stem cells (Johnson et al., 2004). Not one was also Ddx-4 positive. It was possible that the stem cells themselves are not always Ddx4- or Oct4- positive, but if present and active they should give rise to Ddx4-positive, morphologically identifiable oocytes in immature or maturing follicles. 8239 oocyte nuclei were scored in ovaries at all periods of washout – none were IdU positive indicating that none were derived from an adult mitosis (Table 1). Double labelling for IdU and a second germline marker, Oct4 was performed to determine whether any cells in the stem lineage had been missed by the previous analysis. Although all immature oocyte nuclei were found to be Oct4 positive, none were also IdU positive (Figure 3).

**Figure 3.**
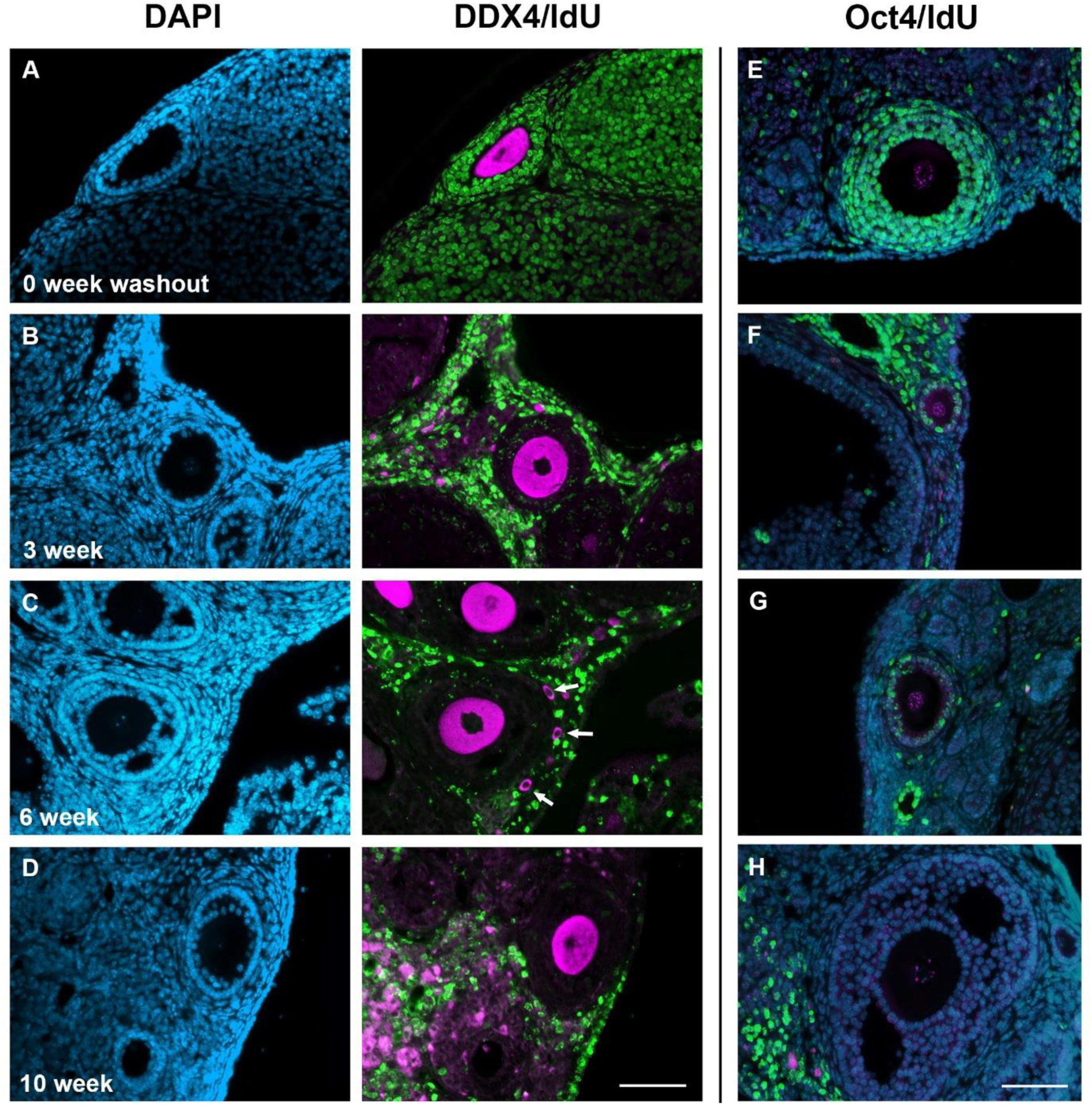
Immunohistochemical double staining of IdU (green) and Ddx4 or Oct4 (magenta) of ovarian follicles at different washout periods. Representative sections of ovaries from young adult mice. All sections stained with DAPI (blue). Ddx4 staining is in the cytoplasm of oocytes. (**A)** Ddx4-positive, IdU-negative oocyte in primary follicle at 0 week washout with IdU-positive granulosa cells. **(B)** IdU-negative oocyte in secondary follicle with Ddx4-positive cytoplasm at 3 weeks washout. IdU has diluted from dividing granulosa cells. **(C, D)**. 6 and 10 weeks washout. Examples are shown of IdU-negative oocytes in follicles starting to become antral. White arrows indicating Ddx4-positive, IdU-negative primordial oocytes. **(E)** Secondary follicle at 0 week washout with Oct4-positive nucleus. Multiple layers of IdU-positive granulosa cells seen but the oocyte is IdU-negative. **(F)** Primary follicle present at 3 weeks washout towards edge of ovary with IdU-negative oocyte nucleus stained positive for Oct4. **(G)** Primary follicle containing oocyte with Oct4-positive nucleus. **(H)** Maturing (tertiary) follicle showing Oct4 positive nucleus of oocyte. All Oct4-positive oocytes were IdU-negative. Scale bar represents 50 μm.

**Table 1:** Quantification of IdU and germline lineage labelling in young adult female mice. Table summarises the analysis of 96 ovaries from 50, 10 week-old C57BL/6 mice, labelled with IdU for 30 days, followed by washout periods of 0-10 weeks. Cells of the ovarian epithelium were counted and scored as IdU-positive or negative. None were Ddx4 positive. Follicles were scored morphologically as primordial, primary, secondary or tertiary and Ddx4-positive oocytes scored as IdU-positive or negative only when the nucleus was visible in the section. None were IdU positive.

| Washout week | No. of Ovaries | EPITHELIUM |  |  | PRIMORDIAL | PRIMARY | SECONDARY | TERTIARY | IdU & DDX4 DOUBLE +ve cells |
| --- | --- | --- | --- | --- | --- | --- | --- | --- | --- |
|  |  | DAPI +ve cells | IDU +ve cells | %IdU | DDX4 +ve cells | DDX4 +ve cells | DDX4 +ve cells | DDX4 +ve cells |  |
| **0** | 38 | 26490 | 2504 | 9.5 | 1885 | 720 | 480 | 192 | **0** |
| **3** | 20 | 15824 | 1965 | 12.4 | 1314 | 650 | 264 | 100 | **0** |
| **6** | 17 | 9577 | 1300 | 13.6 | 900 | 432 | 170 | 80 | **0** |
| **10** | 21 | 6321 | 840 | 13.2 | 674 | 304 | 120 | 62 | **0** |
| **TOTAL** | **96** | **58212** | **6609** |  | **4773** | **2106** | **926** | **434** | **0** |

### Analysis of mitotic activity in the germline of aging female mice

Female mice lose reproductive output from about 6 months old (Virant-Klun, 2015). Therefore, for comparison, 10 ovaries from 5 month old ‘middle-aged’ mice (3 weeks IdU, 3 or 9 weeks washout) and 5 ovaries from post-reproductive 12 month old mice (3 weeks IdU, 2.5 or 9 weeks washout) underwent the label retaining assay using IdU. Their ovaries were recovered for immunohistochemistry for IdU and Ddx4. Epithelial and follicular count was performed as above (Figure 4; Table 2).

**Figure 4.**
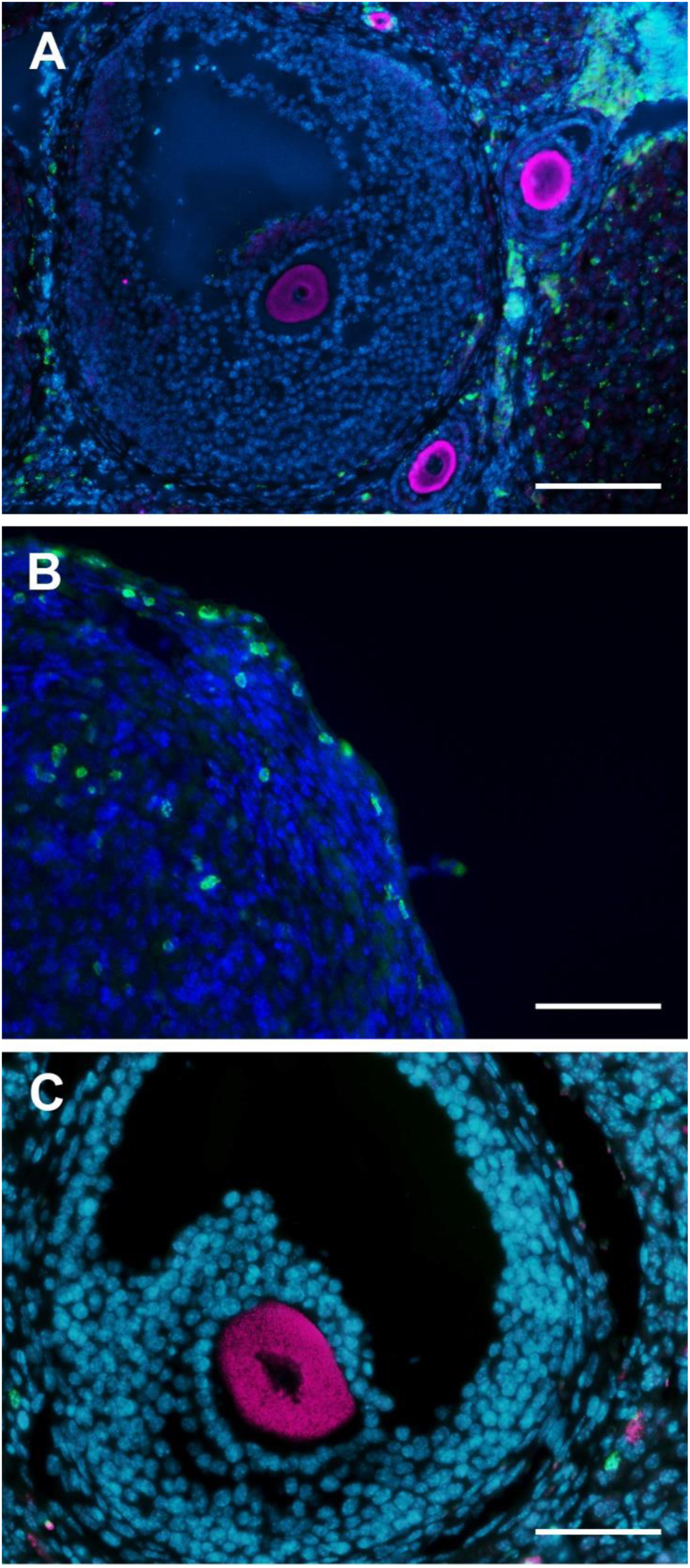
Immunohistochemical double staining of IdU (green) and Ddx4 (magenta) in old mice. All cells stained blue with DAPI. **(A)** Secondary and tertiary follicles from ovaries labelled at 5 months old, positive for Ddx4 but negative for IdU. **(B)** Ovaries from mice with labelling starting at 12 months old were almost devoid of oocytes. IdU label retention observed in epithelium after 9 weeks washout. No Ddx4 labelling. **(C)** Maturing (tertiary) follicle from 12 months ovary stained positive for Ddx4, negative for IdU. Scale bar 50 μm (A), 25 μm (B,C).

**Table 2:** Ageing ovaries. Table summarises the analysis of ovaries of 10, 5 month-old mice and 5, 12 month-old mice labelled with IdU for 21 days followed by washout periods of 2 (1 mouse each timepoint) to 9 weeks (4 mice). Because the previous experiment summarised in table 1 had shown that increasing washout period makes no difference for the oocyte counts, the data for the 9 week and 2 week washout mice are combined here. Cells of the ovarian epithelium were counted and scored as IdU-positive or negative. None were Ddx4 positive. Follicles were scored morphologically as primordial, primary, secondary or tertiary and Ddx4-positive oocytes scored as IdU-positive or negative only when the nucleus was visible in the section. None were IdU positive.

| Age | No. of Ovaries | EPITHELIAL CELLS |  |  | PRIMORDIAL FOLLICLES | PRIMARY FOLLICLES | SECONDARY FOLLICLES | TERTIARY FOLLICLES | IdU & Ddx4 double +ve cells |
| --- | --- | --- | --- | --- | --- | --- | --- | --- | --- |
|  |  | Total number of cells | IDU +ve cells | %IdU | DAPI/DDX4 +ve cells | DAPI/DDX4 +ve cells | DAPI/DDX4 +ve cells | DAPI/DDX4 +ve cells |  |
| **5 months** | 10 | 3250 | 345 | 10.6 | 64 | 52 | 44 | 17 | **0** |
| **12 months** | 5 | 3975 | 523 | 13.1 | 0 | 0 | 0 | 1 | **0** |
| **TOTAL** | **15** | **5225** | **868** |  | **64** | **52** | **44** | **17** | **0** |

In the analysis of the 5 months ovaries, 177 Ddx4-positive oocytes were detected but none were IdU-positive. 345 out of 1250 cortical epithelial cells were IdU positive but none of them were double labelled with Ddx4 (Table 2). As the washout period increased to 9 weeks, follicle morphology and density declined noticeably, but was not quantified.

In the analysis of the 12 months ovaries, there was just one Ddx4-positive oocyte detected in 5 ovaries (Figure 4) and this was not IdU positive. 3975 ovarian cortical epithelial cells were scored and 523 were IdU label-retaining, but none were Ddx4-positive. No IdU/Ddx4 double positive cells were observed in any ovary, consistent with data from younger ovaries.

## DISCUSSION

This study failed to find any oocytes or any other cells double labelled during adulthood for IdU and germline markers, and therefore failed to find any evidence of stem cell activity in the adult ovary at any age. These data are consistent with the multiple studies that have concluded that ovarian germline stem cell activity does not exist (Begum et al., 2008; Kerr et al., 2012; Lui et al., 2007; Zhang et al., 2012; Lei and Spradling, 2013; Zhang et al., 2015; Yoshishara et al., 2023). At the same time, there are many lines of evidence and over 80 publications showing that mitotic ovarian cells can be identified and isolated that, at least in some situations, can produce new oocyte-like cell (Alberico et al., 2022; Guo et al., 2016; Silvestris et al., 2018; Wang et al., 2017; White et al., 2012. Wu et al., 2017; Yang et al., 2018. Reviewed in Woods and Tilly, 2023). Methodological concerns have been raised to explain the apparently contradictory results and reproducibility of stem cell identification (Woods and Tilly, 2015). The suitability of DDX4 protein as a specific female germline marker has been disputed, but defended (Zarate-Garcia et al., 2016; Wang et al., 2017). Since the initial discovery (Johnson et al., 2004), other studies have used short-term thymidine analogue labelling to identify proliferative cells in the female germline, with conflicting positive (e.g. Szotek et al 2008; Guo et al.,2016) or negative (e.g. Lei and Spradling, 2013) results. Reconciling these apparently conflicting lines of evidence requires some assumption that oocyte-regenerating mitotic cells are rare or quiescent in vivo but can be induced to differentiate in vitro, or activated and identified after injuries such as cell dissociation (Telfer and Anderson, 2019). If mitotic germline stem cells are rare, but can be detected after short term (1-3 day) thymidine analogue labelling, then they and their progeny should be obvious after 3 weeks of labelling (as for other slow-cycling stem cells that we studied in Sagga et al., 2018). That we were unable to detect them suggests that they must be quiescent or absent in our mice, but does not preclude activity in other strains or systems including human.

In the current study, ageing was used as a natural, non-invasive route of oocyte depletion but appeared not to lead to activation of any mitotic stem cells. Three key unknowns are how often the germline stem cells might divide (i.e. how long is the required labelling period), whether that changes with age, and how long does an active stem cell take to make a new oocyte in vivo. It is conceivable that the stem cells exist but are quiescent (Pachiarotti et al., 2010), however if none of them are dividing in a 21 - 30 day period, their contribution to replenishing the ovarian stem cell pool was most likely insignificant. Variable washout periods from 0-10 weeks should ensure that oocytes derived from mitoses should be identifiable whether they differentiate and ovulate quickly or slowly, i.e. that the potential to miss them by only looking at one time point is minimised. Our study would also have detected neo-oogenesis, even if the stem cells did not express stem cell or germline markers, or if they produced oocytes that did not express normal germline marker genes, or if the regenerative cells were not true stem cells or if the cells responsible for neo-oogenesis were located outside the ovary. This study also did not rely on expression of transgenic markers. We focused intensively on the ovarian cortical epithelium and found, among many thousands of IdU-retaining cells, none expressing Ddx4 or Oct4 at any age. The analysis would have missed germline stem cells that did not express germline markers. However, oocytes in maturing follicles are easily morphologically recognizable, irrespective of marker expression, so even ‘atypical’ oocytes produced by any sort of mitotic activity in the ovary or anywhere else in the mice should have been detected during the washout periods. We conclude that this mitotic activity probably does not normally exist at biologically relevant levels in vivo, at least in C57BL/6 mice. This is in spite of the fact that it is possible to derive an apparent female germline stem cell line from the ovaries of neonatal C57BL/6 mice (Khosravi-Farsani et al., 2015).

The existence of germline stem cells in postnatal mammalian ovaries remains a controversial issue (Godsen et al., 2004, Byskov et al., 2005; Telfer and Anderson, 2019). As shown by Wu et al. (2022) however, ovarian germline stem cells, even if their activity is ‘only’ experimentally induced, could be fundamental in the treatment of ovarian disease and present a possible therapy for those left infertile due to medical treatments or early menopause.

## Acknowledgements

We thank specialist technical staff at University of Aberdeen Medical Research Facility. ASA was funded by a Ministry of Education in Saudi Arabia PhD studentship. Research work by AFB, SLB, SSK, ST and SLW was funded by the University of Aberdeen as part of their undergraduate studies.

## Authors’ contributions

CMO performed experiments, analysed data, and co-wrote the manuscript. ASA, AFB, SLB, SSK, ST and SLW performed experiments. KK analysed data. JMC conceived and planned the study, performed experiments, and co-wrote the manuscript.

## REFERENCES

Alberico, H., Fleischmann, Z., Bobbitt, T., Takai, Y., Ishihara, O., Seki, H., Anderson, R.A., Telfer, E.E., Woods, D.C. and Tilly, J.L. (2022). Workflow optimization for identification of female germline or oogonial stem cells in human ovarian cortex using single-cell RNA sequence analysis. Stem Cells 40, 523–536.

Altman, J. (1969). Autoradiographic and histological studies of postnatal neurogenesis. IV. Cell proliferation and migration in the anterior forebrain, with special reference to persisting neurogenesis in the olfactory bulb. J Comp Neurol 137, 433–457.

Begum, S., Papaioannou, V.E. & Gosden, R.G. (2008). The oocyte population is not renewed in transplanted or irradiated adult ovaries. Human Reproduction, 23, 2326–2330.

Byskov, A.G., Faddy, M.J., Lemmen, J.G. & Andersen, C.Y. (2005). Eggs forever? Differentiation 73, pp. 438–446.

Carrera, P., Johnstone, O., Nakamura, A., Casanova, J., Jäckle, H., Lasko, P. (2000). VASA mediates translation through interaction with a Drosophila ylF2 homolog. Molecular Cell 5, 181 – 187.

Cotsarelis, G., Cheng, S.Z., Dong, G., Sun, T.T., Lavker, R.M., 1989. Existence of slow-cycling limbal epithelial basal cells that can be preferentially stimulated to proliferate: implications on epithelial stem cells. Cell 57, 201–209.

Erez, R. (2000). The function and regulation of vasa-like genes in germ-cell development. Genome Biol 1, 1017.1 – 1017.6.

Gosden, R.G. (2004). Germline stem cells in the postnatal ovary: is the ovary more like a testis? Hum Reprod Update 10, 193–195.

Grive, K. J., & Freiman, R. N. (2015). The developmental origins of the mammalian ovarian reserve. Development 142, 2554–63.

Guo, K., Li, C.-H., Wang, X.-Y., He, D.-J. and Zheng, P. (2016). Germ stem cells are active in postnatal mouse ovary under physiological conditions. Mol. Hum. Reprod. 22, 316–328.

Izadyar, F., Pau, F., Marh, J., Slepko, N., Wang, T., Gonzalez, R., Ramos, T., Howerton, K., Sayre, C., Silva, F. (2008). Generation of multipotent cell lines from a distinct population of male germ line stem cells. Reproduction 135, 771–784.

Johnson, J., Canning, J., Kaneko, T., Pru, J.K. & Tilly, J.L. (2004). Germline stem cells and follicular renewal in the postnatal mammalian ovary. Nature 428, 145–150.

Johnson, J., Bagley, J., Skaznik-Wikiel, M., Lee, H.J., Adams, G.B., Niikura, Y., Tschudy, K.S., Tilly, J.C., Cortes, M.L., Forkert, R., Spitzer, T., Iacomini, J., Scadden, D.T., Tilly, J.L. (2005). Oocyte generation in adult mammalian ovaries by putative germ cells in bone marrow and peripheral blood. Cell 122, 303–315.

Kerr, J., Duckett, R., Myers, M., Britt, K., Mladenovska, T. and Findlay, J. (2006). Quantification of healthy follicles in the neonatal and adult mouse ovary: evidence for maintenance of primordial follicle supply. Reproduction 132, pp.95–109.

Kerr, J.B., Brogan, L., Myers, M., Hutt, K.J., Mladenovska, T., Ricardo, S., Hamza, K., Scott, C.L., Strasser, A. and Findlay, J.K. (2012). The primordial follicle reserve is not renewed after chemical or γ-irradiation mediated depletion. Reproduction, 143, 469–476.

Khosravi-Farsani, S., Amidi, F., Roudkenar, M.H. and Sobhani, A. (2015) Isolation and enrichment of mouse female germ line stem cells. Cell J. 16, 406–415.

Lei, L and Spradling, A.C. 2013. Female mice lack adult germ-line stem cells but sustain oogenesis using stable primordial follicles. Proc Natl Acad Sci USA 21, 8585–8590.

Liu, Y., Wu, C., Lyu, Q., Yang, D., Albertini, D., Keefe, D. and Liu, L. (2007). Germline stem cells and neo-oogenesis in the adult human ovary. Dev Biol 306, 112–120.

Niikura, Y. Niikura, T. and Tilly, J.L. (2009), Aged mouse ovaries possess rare premeiotic germ cells that can generate oocytes following transplantation into a young host environment. Aging 1, 971–978.

Niikura, Y., Niikura, T., Wang, N., Satirapod, C. and Tilly, J.L. (2010). Systemic signals in aged males exert potent rejuvenating effects on the ovarian follicle reserve in mammalian females. Aging 2, 999–1003.

Pacchiarotti, J., Maki, C., Ramos, T., Marh. J., Howerton, K., Wong, J., Pham, J., Anorve, S., Chow, Y.C., Izadyar, F. (2010). Differentiation potential of germ line stem cells derived from the postnatal mouse ovary. Differentiation, 79, 159–170.

Park, E.S. and Tilly, J.L. (2015). Use of DEAD-box polypeptide 4 (Ddx4) gene promoter-driven fluorescent reporter mice to identify mitotically active germ cells in postnatal mouse ovaries. Mol. Hum. Reprod. 21, 58–65.

Sagga, N., Kuffová L., Vargesson, N., Erskine, L. and Collinson, J. M. 2018. Limbal epithelial stem cell activity and corneal epithelial cell cycle parameters in adult and aging mice. Stem Cell Res. 33: 185–198.

Silvestris, E., Cafforio, P., D’Oronzo, S., Felici, C., Silvestris, F. and Loverro, G. (2018) In vitro differentiation of human oocyte-like cells from oogonial stem cells: Single-cell isolation and molecular characterization. Hum. Reprod. 33, 464–473.

Szotek P, Chang H, Brennand K et al. (2008) Normal ovarian surface epithelial label-retaining cells exhibit stem/progenitor cell characteristics. Proc Natl Acad Sci (USA) 105, 12468–12473.

Telfer EE, Anderson RA 2019. The existence and potential of germline stem cells in the adult mammalian ovary. Climacteric 22, 22–26.

Tilly, J.L. & Telfer, E.E. (2009). Purification of germline stem cells from adult mammalian ovaries: a step closer towards control of the female biological clock? Mol Hum Reprod 15, 393–398.

Virant-Klun I. (2015). Postnatal oogenesis in humans: a review of recent findings. Stem Cells Cloning 8, 49–60.

Wang, N., Satirapod, C, Ohguchi, Y., Park, E.-S., Woods, D.C. and Tilly, J.L. (2017). Genetic studies in mice directly link oocytes produced during adulthood to ovarian function and natural fertility. Sci. Rep. 7, 10011.

White, Y.A., Woods, D.C., Takai, Y., Ishihara, O., Seki, H. and Tilly, J.L. (2012). Oocyte formation by mitotically active germ cells purified from ovaries of reproductive-age women. Nature Medicine, 18, 413–421.

Woods, D.C. and Tilly, J.L. (2013). Isolation, characterization and propagation of mitotically active germ cells from adult mouse and human ovaries. Nat. Protoc. 8, 966–988.

Woods, D.C. and Tilly, J.L. 2015. Reply to adult human and mouse ovaries lack DDX4-expressing functional oogonial stem cells. Nat. Med. 21, 1118–1121.

Woods, D. C. and Tilly, J. L. (2023) Revisiting Claims of the Continued Absence of Functional Germline Stem Cells in Adult Ovaries. Stem Cells 41, 200–204. 10.1093/stmcls/sxac083.

Wu, C., Xu, B., Li, X., Ma, W., Zhang, P. and Wu, J. (2017). Tracing and characterizing the development of transplanted female germline stem cells in vivo. Mol. Ther. 25, 1408–1419

Wu, M., Lu, Z. Zhu, Q., Ma, L., Xue, L., Li, Y., Zhou, S., Yan, W., Ye, W., Zhang, J., Luo, A. and Wang, S. (2022). DDX04^+^ Stem Cells in the Ovaries of Postmenopausal Women: Existence and Differentiation Potential. Stem Cells, 40, 88–101. 10.1093/stmcls/sxab002.

Yang, H., Yao, X., Tang, F., Wei, Y., Hua, J., Peng, S. (2018). Characterization of female germline stem cells from adult mouse ovaries and the role of rapamycin on them. Cytotechnol. 70, 843–854.

Yoshihara, M., Wagner, M., Damdimopoulos, A., Zhao, C., Petropoulos, S., Katayama, S., Kere, J., Lanner, F. and Damdimopoulou, P. (2023). The Continued Absence of Functional Germline Stem Cells in Adult Ovaries, Stem Cells, 41, 105–110. 10.1093/stmcls/sxac070.

Zarate-Garcia, L., Lane, S. I., Merriman, J. A., & Jones, K. T. (2016). FACS-sorted putative oogonial stem cells from the ovary are neither DDX4-positive nor germ cells. Scientific Rep 6, 27991. doi:10.1038/srep27991

Zhang, H., Zheng, W., Shen, Y., Adhikari, D., Ueno, H., & Liu, K. (2012). Experimental evidence showing that no mitotically active female germline progenitors exist in postnatal mouse ovaries. Proc Natl Acad Sci (USA). 109, 12580 – 12585.

Zhang, H., Panula, S., Petropoulos, S., Edsgärd, D., Busayavalasa, K., Liu, L., Li, X., Risal, S., Shen, Y., Shao, J., Liu, M., Li, S., Zhang, D., Zhang, X., Gerner, R. R., Sheikhi, M., Damdimopoulou, P., Sandberg, R., Douagi, I., Gustafsson, J. Å., Liu, L., Lanner, F., Hovatta, O., & Liu, K. (2015). Adult human and mouse ovaries lack DDX4-expressing functional oogonial stem cells. Nature Medicine. 21, 1116 – 1118.

Zou, K., Yuan, Z., Yang, Z., Luo, H., Sun, K., Zhou, L., Xiang, J., Shi, L., Yu, Q., Zhang, Y., Hou, R., Wu, J. (2009). Production of offspring from a germline stem cell line derived from neonatal ovaries. Nature Cell Biol 11, 631–636.

Zou, K., Hou, L., Sun, K., Xie, W., Wu, J. (2011). Improved efficiency of female germline stem cell purification using Fragilis-based magnetic bead sorting. Stem Cells Dev., 20, 2197–2204.

Zuckerman, S. (1951). The number of oocytes in the mouse ovary. Recent Prog Hormone Res 6, 63– 108.

